# MAESTRO uncovers tandem paralog dynamics as a core driver of fungal stress adaptation

**DOI:** 10.1101/2025.08.29.673034

**Authors:** Ping-Hung Hsieh, Yusuke Sasaki, Chu-I Yang, Zong-Yen Wu, Andrei S. Steindorff, Sajeet Haridas, Jing Ke, Zia Fatma, Zhiying Zhao, Dana A. Opulente, Siwen Deng, Chris Todd Hittinger, Igor V. Grigoriev, Bruce Dien, Huimin Zhao, Yi-Pei Li, Yasuo Yoshikuni

## Abstract

Understanding how fungi adapt to diverse stresses is critical for mitigating emerging drug resistance and harnessing their robustness for biotechnology. *Pichia kudriavzevii* is a stress-tolerant and intrinsically drug-resistant yeast, with dual industrial and clinical importance. We analyzed 170 strains using MAESTRO, a machine-learning-assisted GWAS pipeline optimized for small cohorts. MAESTRO identified biologically meaningful features and revealed that copy number variation (CNV) of tandem paralogs (TPs) is a core mechanism of multi-stress adaptation. TPs were recurrently linked to tolerance of industrial inhibitors (HMF, phenolics, heat) and antifungal drugs (fluconazole, azoles), and deletion of the TP pair gene4260/gene4261 confirmed pleiotropic effects across stresses. These findings support a TP CNV model where recombination-driven TP CNVs and gene fusions enable rapid stress adaptation. Importantly, our results suggest that antifungal resistance can arise through co-option of mechanisms originally evolved for environmental stressors, raising a One Health concern about the environmental origins of drug-resistant pathogens.

## INTRODUCTION

Fungi can be our foes or our friends. As pathogens, these eukaryotic organisms cause more than a billion human infections each year worldwide, and 1.6 million people die annually from invasive fungal diseases^1,2^. In agriculture, hundreds of fungal diseases affect important crops, causing farmers to lose 10% to 23% of their crops every year before harvest and another 10% to 20% post-harvest^3,4^. Climate change is thought to worsen the problem by expanding habitats favorable to pathogens (e.g., wheat-stem rust infections)^3^ and activating dormant pathogens (e.g., *Candida auris*)^5^. As our allies, many fungal strains are used to produce foods and beverages, while some serve as engineering hosts for production of diverse bio-based products^6^. Recent reports predict that the bioeconomy will grow to from $2 trillion to $4 trillion by 2040, and fungi are expected to play important roles in this growth as chassis for production of pharmaceuticals, consumer products, renewable chemicals, and biofuels^7,8^. Clearly, understanding fungal evolution is critical to both managing their risks on human health and harnessing their benefits on bioeconomy^4,6,9–12^

Unlike bacteria, which can acquire genes rapidly through horizontal gene transfer (HGT), fungi rarely undergo HGT^4^. For example, genomic analysis of budding yeasts shows that only perhaps 0.04% to 0.06% of annotated genes are acquired through HGT^13^. Instead, the predominant mechanisms underlying fungal evolution are thought to be asexual mutations and sexual reproduction^4,14^. Short variants, such as single nucleotide polymorphisms (SNPs) and small insertions and deletions (indels), had been believed to be major contributors to large genotypic and phenotypic diversity. However, recent advances in sequencing technologies are revealing that structural variations (SVs) such as polyploidy^15^, aneuploidy^16,17^, loss of heterozygosity (LOH)^18–20^, and gene copy number variations (CNVs)^21,22^ may be equally or even more crucial than short variants in fungal adaptive evolution^21,23^. Nonetheless, the relationship between SVs and fungal adaptive evolution is still underexplored. Improving our understanding of fungal evolution driven by both short variants and SVs could help us develop strategies for detecting high-risk pathogens, developing effective counteragents, and designing and building fungal strains more suitable for industrial uses.

For this study, we selected *Pichia kudriavzevii* (a.k.a. *Issatchenckia orientalis, Candida krusei*) as a model^24^. *P. kudriavzevii* can be harmful or helpful to humans. On one hand, it is a multidrug-resistant opportunistic pathogen that causes invasive fungal diseases and is on the WHO’s list of 19 priority fungal pathogens^1,2^. On the other hand, *P. kudriavzevii* is used to produce foods and beverages (e.g., chocolate, coffee, liquor), and some strains are even generally recognized as safe (GRAS) by the FDA^25^. Additionally, *P. kudriavzevii* has extraordinary ability to tolerate multiple stressors and is recognized as an emerging industrial host^26–32^. Studies have demonstrated that low-pH fermentation by *P. kudriavzevii* to produce organic acids, which are building blocks for various plastics and polymers, could reduce the cost and environmental footprint by 23% relative to the best-case scenario using the conventional process^33^. Several companies and academic research groups are developing *P. kudriavzevii* as a chassis for production of diverse bioproducts^30–32,34–38^. Despite the importance of this species, most genomics studies have been limited to small sample sizes and have primarily focused on adaptive genetic drivers that are already known, such as known genes or homologs from closely related species^24^.

To more comprehensively identify the major drivers of adaptive evolution in P. kudriavzevii, we performed population genomics and phenomics studies, followed by a genome-wide association study (GWAS) using both short variants and CNVs. For the GWAS, we developed a computational pipeline, “machine-learning-assisted engineering of stress-tolerant rational optimization (MAESTRO).” Using a combination of machine-learning methods with cross-validation and optimization, MAESTRO identified sets of polygenic features that effectively correlate genotypes (including both short variants and CNVs) with phenotypes. Together with experimental validation, these studies revealed tandem paralogs and CNV dynamics as key mechanisms of fungal stress adaptation, while also demonstrating MAESTRO as a practical framework for extracting biologically meaningful associations in systems where sample size is limited.

## RESULTS

### P. kudriavzevii *strain collection*

We collected 170 strains from around the globe (Fig. S1A and Table S1) and classified them into seven groups according to where they were found (their origin of isolation): on edibles (food and drinks); on plants or animals (including insects); in the general environment, industrial settings, or clinical settings; or from unknown places (Fig. S1B).

### *Population genomics of* P. kudriavzevii *strain collection*

We sequenced, assembled, and annotated the genomes of 170 strains (Fig. S2, see Methods), identifying 2,719 single-copy orthologs, which we used to construct a phylogenetic tree (Fig. 1). We also mapped the sequencing reads from these strains to the reference genome of strain CBS573^24^ (also resequenced and denoted as IO11) and identified 308,759 biallelic short variants (SNPs and indels)^39^ (Table S2). Most of the short variants were in intergenic regions and occurred at a low frequency, with 70% showing a minor allele frequency of below 5% (Fig. S3). An ADMIXTURE analysis based on the filtered short variants indicated that the strains in our population likely originated from 14 ancestral groups, with 27% classified as mosaic strains, defined as strains consisting of less than 80% of a single ancestral population (Fig. 1, Table S3). Using the same filtered short variants, a principal component analysis (PCA) revealed three main groups, which we organized into populations 1–9, 10 and 11, and 12–14; these groups were interconnected by mosaic strains (Fig. S4). Notably, the results from phylogenetic, ADMIXTURE, and PCA analyses were largely consistent.

**Figure 1.**
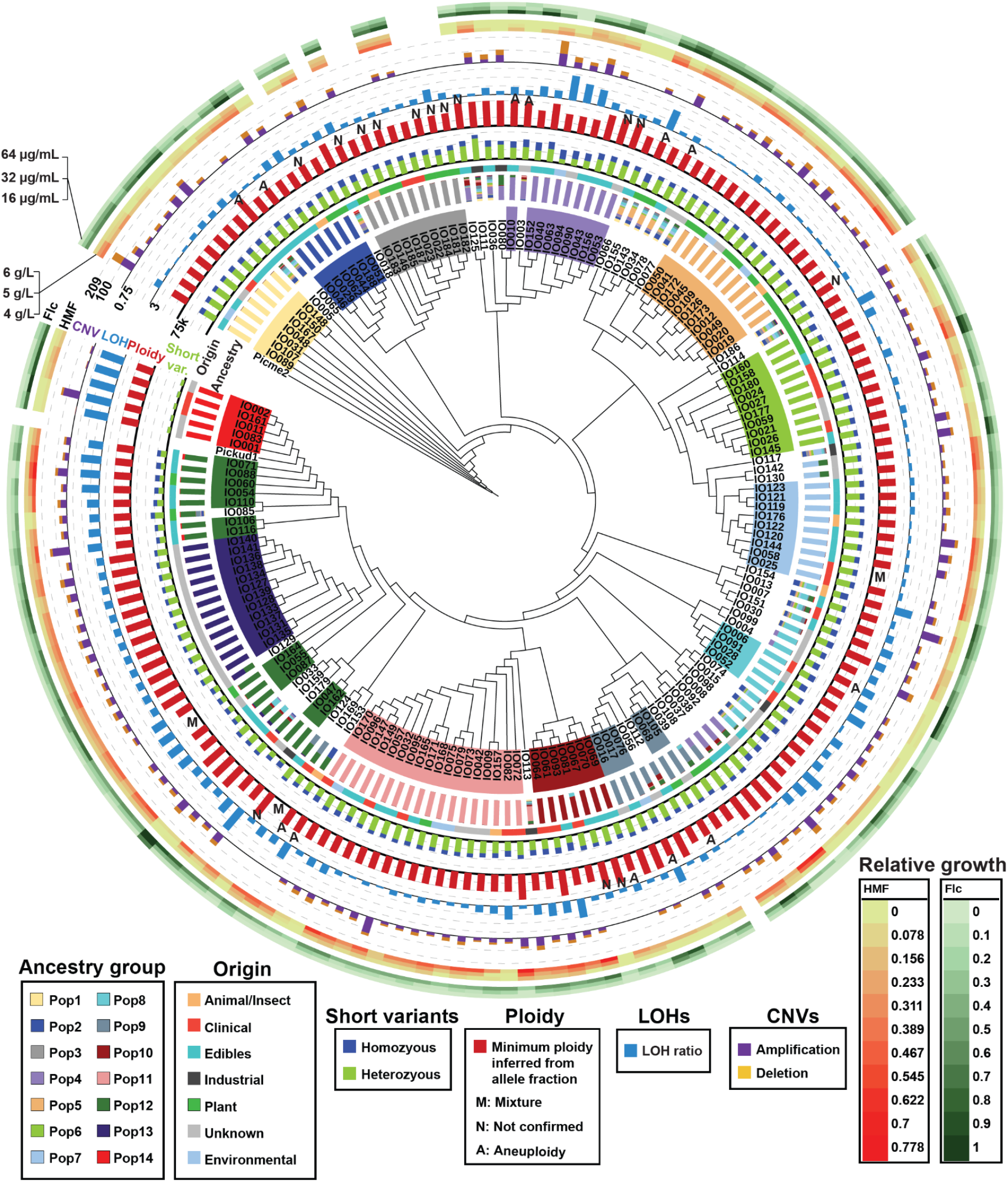
Population genomics study of 170 *P. kudriavzevii* strains. (A) This dendrogram presents the maximum-likelihood phylogenetic tree of 170 strains. It was constructed based on 2,719 single-copy orthogroups. *Pichia membranifaciens* (denoted as Picme2) was used as an outgroup. *Pichia kudriavzevii* strain CBS573 (denoted as Pickud1) was used as a reference genome (accession: PRJNA434433), which was assembled using a PacBio platform. We also resequenced the CBS573 strain using an Illumina platform, and this is denoted as IO11. The colors used to highlight each strain name and those in the “Ancestry” column reflect the ancestral populations determined by ADMIXTURE. Mosaic strains, defined as strains consisting of less than 80% of a single ancestral population, are not highlighted. The “Origin” column identifies the habitats from which each strain was isolated (Table S1). The “short variant counts” column indicates the number of homozygous and heterozygous short variants detected by GATK (Table S3). The “Ploidy” column shows the inferred minimum ploidy of strains, derived from the sequencing datasets (Table S4). **M**: a mixed population, **N**: not confirmed by flow cytometry, and **A**: segmental aneuploidy (Table S5). The “LOH” column presents the ratio of the genome for each genome exhibiting LOH (Table S6). The “CNVs” column displays gene amplification and deletion (reduced copy number and loss) events for each strain (Table S7). Only the non-mixture strains with confirmed ploidy and no segmental aneuploidy were used for CNV analysis. The heatmaps exhibit normalized growth (colony sizes), calculated as the ratio of growth under various concentrations of fluconazole (Flc) and 5-hydroxymethylfurfural (HMF) to the growth observed under control conditions (Tables S8 and S9).

We also explored potential correlations between these strains’ evolutionary relationships and isolation origins using a Sankey plot (Fig. S5). However, we found no correlation, suggesting that *P. kudriavzevii* strains are likely circulating in diverse ecosystems^24^. As an example, our strains IO181 through IO187 had been isolated from trees in Madison, Wisconsin, USA, (Table S1) and were clustered together in the same clade, except for the mosaic IO186 (Fig. 1). However, this clade also included three other isolates from clinical and animal sources: IO23, IO24, and IO146 (Fig. 1).

Investigating ploidy variations using methods based on flow cytometry and allele fraction (Fig. 1, S6, and Table S4), we cross-validated 119 diploid and 34 triploid strains. Because of the high flocculence of the remaining 17 strains, we used only the allele-fraction-based approach to infer their minimum ploidy. Surprisingly, no haploid strain was identified in our collection (Table S4). We also used the sequencing data to identify a total of 17 segmental aneuploidies across 13 strains, ranging from 25 kb to entire chromosomes (Fig. S7, S8, and Table S5). Chromosome 4 exhibited the highest number of segmental aneuploidy events (Fig. S8). Analyzing loss of heterozygosity (LOH), we identified substantial variation across strains; the maximum size of continuous LOHs ranged from 0.055 Mb to 0.28 Mb, and maximum cumulative LOH ranged from 0.28 Mb to 7.5 Mb (Fig. 1 and Table S6). Consistent with previous studies,^21,24^ LOH was more frequent on chromosome arms than on centromeres (Fig. S9).

We used a pangenome analysis pipeline^21^ to identify open reading frames (ORFs) not found in the reference strain CBS573. This analysis identified only five non-reference ORFs and 79 gene truncations or loss events, suggesting that the *P. kudriavzevii* population has a large set of core genes (> 98%). Therefore, we focused our CNV analysis on the 4,757 reference genes and excluded aneuploid strains (Fig. 1, S10, and Table S7). In at least one strain, 13.6% to 26.1% of genes had been amplified relative to the reference strain in each chromosome. In contrast, 8.5% to 12.2% of genes had been truncated or lost (Fig. S10A). However, these CNVs were rare at the population level; about 30% of gene amplifications and 40% of deletions were specific to a single strain (Fig. S10B and C). Consistently, the mean probability of gene amplification was between 0.7% and 1.6%, and the mean probability of gene deletion was between 0.2% and 1% per chromosome (Fig. S10D). This phenomenon aligns with the low population frequency of short variants, suggesting a recent divergence among strains or the presence of significant selection pressure on CNVs. Interestingly, we found that non-clinical strains have more genes with CNVs than the clinical strains (Mann–Whitney U test, p < 0.01, Fig. S10E).

### Population phenomics of *P. kudriavzevii strain collection*

To investigate the fitness of each strain under diverse stress conditions, we developed a high-throughput (HTP) solid-agar-based growth assay with an Echo acoustic liquid handling system (Fig. S11). This system allowed us to manipulate cell cultures at the nanoliter scale and to eject the cultures onto agar plates at a density of 384 distinguishable spots. We selected 57 stress conditions (32 antifungal and 25 industrially relevant) for testing in the HTP fitness study and collected 9,003 datapoints (Fig. S11–S13). For all these datapoints, we confirmed high reproducibility among 3–6 technical replicates and between two biological replicates (median Spearman correlation coefficient ρ = 0.84; a linear model, coefficient of determination R^2^ = 0.92, p < 10^-3^) after filtering out the datapoints with high variation. We calculated fitness scores (Z-scores) for each strain under the different stress conditions and created a fitness profile matrix as a heatmap (Fig. 2, Tables S8 and S9). We adopted a tree-based strain classification procedure and categorized strains into several classes using a hierarchical clustering analysis. This analysis classified the whole set of strains into three major populations (A–C) and six subpopulations (A, B1-2, C1–3) (Fig. 2 and S14).

**Figure 2.**
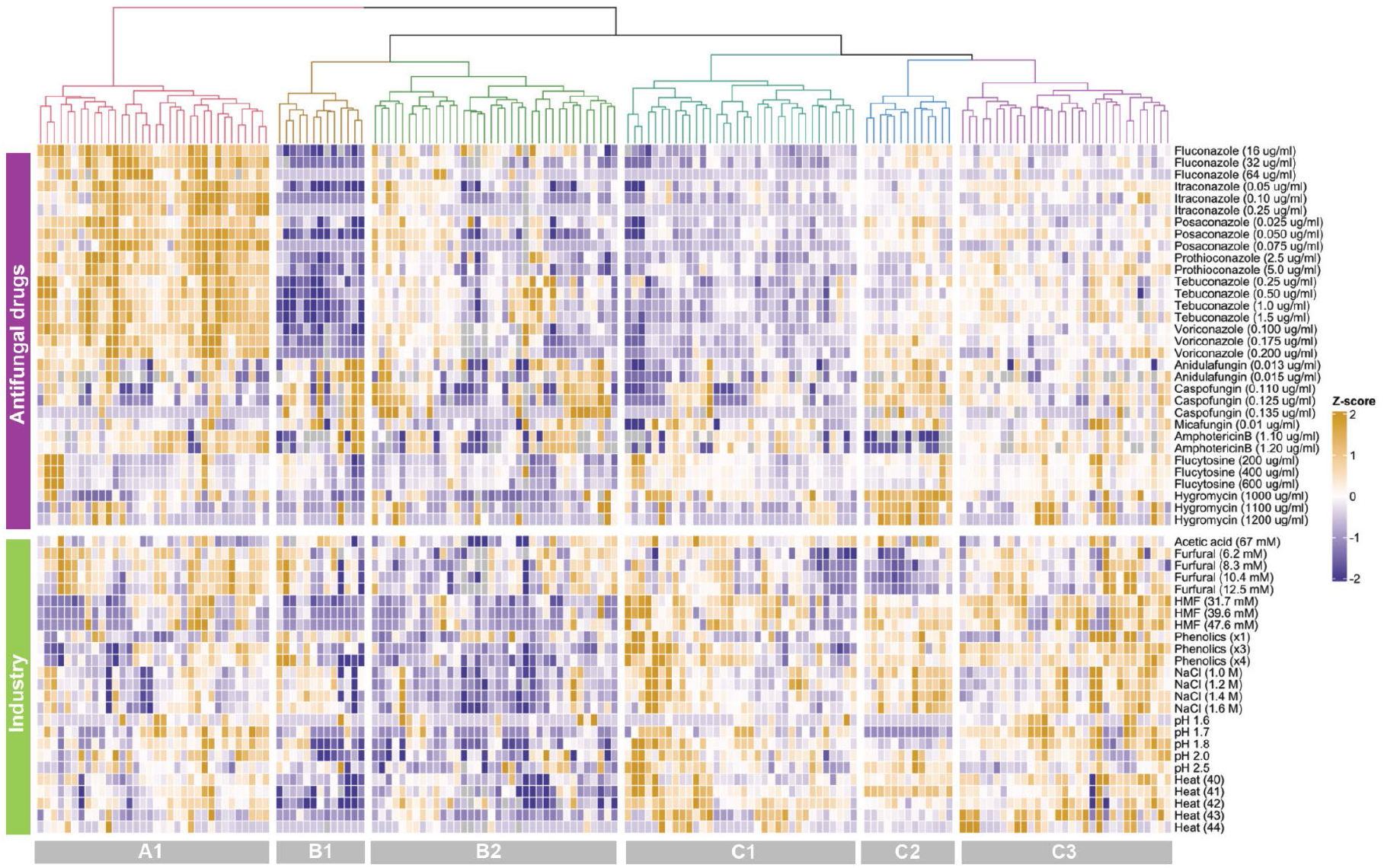
Fitness landscape of the *P. kudriavzevii* population and relationships to ecological origins. (A) The results for the population phenomics study of 161 *P. kudriavzevii* strains are summarized as a heatmap. For each condition, the fitness diversity among this population was normalized using Z-scores. The normalized fitness scores are represented with the scale bar next to the plot. Missing values (no data) are shown in gray. A hierarchical clustering analysis was used to group the *P. kudriavzevii* population based on normalized fitness scores, which resulted in three populations and six subpopulations (A1, B1-2, and C1–3).

We subsequently evaluated phenotypic variations among subpopulations using permutation analysis of variance (PERMANOVA) and confirmed that these subpopulations were significantly different from each other (pseudo-F = 7.33, p-value < 10^-3^). We visualized the relationships among strains and the populations by embedding them in a low-dimensional space using several approaches, supporting the validity of the hierarchical clustering result (Fig. S15). The traits of each population under antifungal-resistance and industrially relevant conditions were summarized using violin plots (Fig. S16). We also investigated the relationship between their phenotypes and their origins of isolation (Fig. S17 and S18) but found no obvious correlations.

### Population-wide association study

We performed this study for a few traits such as resistance to fluconazole (an antifungal [clinical application]) and tolerance of 5-hydroxymethyl furfural (HMF) (one of inhibitors in lignocellulosic hydrolysate^40^ [industrial application]) as examples. We initially studied whether there were any obvious correlations between phenotypes and ADMIXTURE groups (Fig. S19), finding no statistically significant correlations. Additionally, we examined the effect of ploidy variations on these traits, finding no correlation between them either. However, we found that diploids grew slightly better than triploids did under the reference condition (Fig. S20). Investigating the effect of LOH, we found that cumulative LOH has a weak negative correlation with HMF tolerance (p-value < 0.05) and a weak positive correlation with fluconazole resistance (not significant) (Fig. S21). We continued the investigation at the level of clades. We mapped fitness scores of each strain under different stress conditions to the phylogenetic tree (Fig. 1). This analysis showed that two evolutionarily closely related strains could have greatly different phenotypes. For example, strains IO24 and IO27 are closely related, but IO27 showed significantly higher resistance to fluconazole and tolerance of HMF than did IO24. Across this phylogenetic tree, we identified other similar examples (e.g., IO033 and IO159, IO085 and IO110). These results suggest that relatively small numbers of GVs are sufficient for phenotypic diversification. A deeper analysis is needed to determine the contribution of individual GVs to fungal adaptive evolution.

### ML-assisted GWAS using MAESTRO

We first used the linear mixed model^41^ to specifically analyze short variants associated with fluconazole resistance and HMF tolerance. However, none of the short variants met the Bonferroni significance threshold. This limitation is likely attributable to the small sample size of our collection and the polygenic nature of the stress-tolerance phenotypes^42,43^. Nevertheless, the short variants whose correlations with stress response had highest significance were experimentally validated. While overexpression of gene305 increased HMF tolerance, deletion of gene1682(MSN4) and gene1677(BYE1) slightly increased fluconazole resistance (Fig. S22 and Table S10).

Previous studies have demonstrated that machine learning approaches can capture complex, non-linear genetic effects that might be overlooked by traditional GWAS^44–46^. As a complementary strategy, we developed MAESTRO, an ML-assisted GWAS pipeline that integrates feature selection and selection-frequency-based evaluation to identify stable genetic features underlying genotype–phenotype associations. (Fig. 3A). Because our dataset for GWAS is limited (160 samples), holding out a separate test set would easily exclude unique samples that contain critical genotype–phenotype signals. Indeed, we observed that even by removing even 5% of samples could prevent us from identifying genetic variants from known causal genes during internal validation, making the conventional held-out strategy not feasible. In finite and high-dimensional datasets, stable feature selection via repeated resampling has been shown to increase the likelihood of identifying truly relevant features by reducing false discoveries and making the selection outcome less sensitive to the choice of regularization^47^. Therefore, we designed our pipeline to integrate stable feature selection through 10 rounds of 10-fold nested cross-validation (CV) with recursive feature elimination (RFE) based on linear regression, which repeatedly re-samples the data to identify consistently selected genetic variants associated with the traits. By aggregating results from 1,000 feature selection iterations, we obtained selection frequencies for each feature, where a higher frequency indicates a greater likelihood of representing a truly relevant variant.

**Figure 3.**
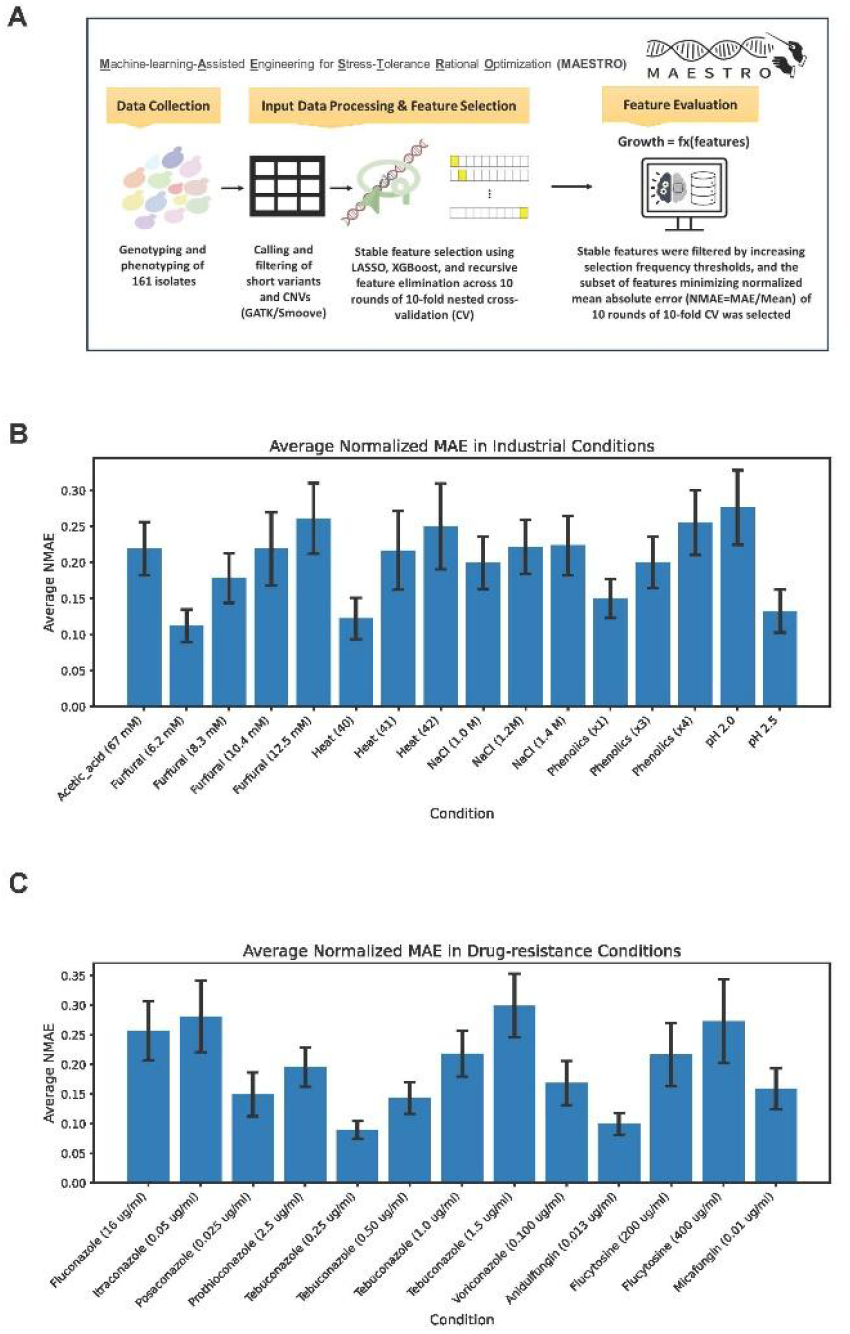
MAESTRO pipeline and prediction performance across stress conditions. (A) Overview of the MAESTRO pipeline. Genotypes and phenotypes were processed to call SNPs/Indels and CNVs, followed by stable feature selection using nested cross-validation and recursive feature elimination. Selection frequencies were aggregated across 1,000 iterations, and an additional CV step was used to determine the optimal cutoff for final feature sets. (B–C) Prediction accuracy of MAESTRO-selected CNVs across 16 industrial and 13 drug-resistance conditions. Bars show average NMAE across folds (error bars: standard deviation). Conditions such as furfural, heat (40 °C), tebuconazole, and posaconazole achieved particularly low error, highlighting stress contexts where CNV-based predictions were most reliable.

As the empirical stability cutoff (0.6–0.9) was too stringent for our limited dataset, we implemented an additional 10-rounds of 10-fold CV analysis to determine the optimal frequency threshold for the final feature set. We systematically evaluated all unique selection-frequency cutoffs from 0.1 to the maximum observed frequency. For each threshold, features with mean selection frequency greater than or equal to the cutoff were retained, and linear regression models were trained across the 10 folds of cross-validation. Performance was evaluated using normalized mean absolute error (NMAE = MAE divided by the mean of the observed values), which enables fair comparison across stress conditions with different phenotype ranges. The cutoff that minimized the average NMAE across folds was then selected to define the final feature set for each stress condition. Although this design cannot entirely eliminate the risk of data leakage, it offers a balance route to maximize the utility of limited data while minimizing overfitting.

As a sanity check, we applied the MAESTRO pipeline to fluconazole resistance using either SNPs/Indels or gene CNVs as input features and then examined whether the selected variants included known benchmark genes such as ERG11 (the fluconazole drug target) and ABC1 (an ABC transporter induced by azoles)^48^. When using SNPs/Indels for the 16 µg/mL fluconazole condition, 9 of the top 10 most frequently selected features were located in intergenic regions. Because of linkage disequilibrium (LD), it is challenging to reliably identify which of the nearby genes may be causal, and our simple inspection of the closest genes to these SNPs/Indels did not reveal known fluconazole resistance targets. In contrast, when gene CNVs were used as input, the most frequently selected feature was the CNV of the known fluconazole resistance gene ABC1p (gene4260), which appeared at high selection frequencies (0.694 at 16 µg/mL and 0.697 at 32 µg/mL out of 1000 times of feature selection). While previous studies have shown that ABC1 is overexpressed in azole-resistant strains^49^, our results newly demonstrate that CNV of ABC1 may also contributes to resistance in *P. kudriavzevii*. Because of the difficulty in assigning causal genes from SNPs/Indels under LD constraints, and the clear recovery of a known resistance gene from gene CNV analysis, we focused our downstream analyses only on gene CNVs.

Using this integrated pipeline, we defined reliable prediction as either NMAE < 0.3 or R^2^ > 0.3. Across the 57 tested stress conditions, MAESTRO-selected CNVs achieved NMAE < 0.3 in 16 industrial stresses and 13 drug-resistance conditions (Figs. 3B, 3C). Conditions such as 6.2 mM furfural, heat at 40 °C, and 0.25 µg/mL tebuconazole showed particularly low NMAE around 0.1, highlighting cases where CNV-based predictions were more reliable. In addition, four other conditions (31.7 mM HMF, 39.6 mM HMF, 0.050 µg/mL posaconazole, and pH 1.8) reached R^2^ > 0.3 while they can’t meet the NMAE < 0.3 threshold. Notably, although CNV-based predictions for HMF did not reach the low NMAE standard, the most frequently selected feature was the CNV of a homolog of the *Saccharomyces cerevisiae* alcohol dehydrogenase ADH6p that detoxifies HMF by converting it to 5-hydroxymethylfurfuryl alcohol^50^. This suggests that selected features from our pipeline can provide biologically meaningful insights even when overall prediction accuracy is limited.

### Tandem paralogs are consistently associated with multiple stress conditions

The identification of ABC1 in fluconazole resistance and an ADH6 homolog in HMF tolerance suggests that MAESTRO-selected CNV features can capture biologically meaningful targets, rather than serving only as predictive variables. Further exploring features across conditions, we surprisingly found that tandem paralogs (TPs) were recurrently selected across diverse stress conditions (Fig. 4A). This enrichment was not observed for non-tandem paralogs, indicating that the enrichment pattern is unlikely due to alignment artifacts among homologous sequences (Fig. 4A). TPs are well recognized as hotspots for non-allelic homologous recombination (NAHR)^51^, which facilitates copy number amplification or reduction and can also generate new functions through in-frame gene fusion. Supporting this view, we found that TPs had significantly higher CNV frequencies across strains than either non-paralog genes or non-tandem paralogs (Fig. 4B). Such structural plasticity may confer selective advantages under changing environments. At the functional level, gene ontology (GO) enrichment analysis showed strong enrichments of transporters and other membrane proteins among TPs (Fig. 4C). InterProScan annotations further revealed that TPs are significantly enriched for transporters (52 of 108 TPs, Fisher’s exact test p-value < 0.01, Fig. S23) compared with the whole genome (233 of 4,757 genes). Together, these structural and functional enrichments suggest that TPs play important roles in stress adaptation. To pinpoint specific candidates, we next examined which TPs were recurrently selected across stress conditions (Fig. 4D). Our analysis revealed candidate genes with potential pleiotropic effects on stress responses. Surprisingly, the most recurrent TP across conditions was gene4260, a known fluconazole-resistant ABC transporter, which was also associated with tolerance to phenolics, HMF, heat, and low-pH (Fig. 4D).

**Figure 4.**
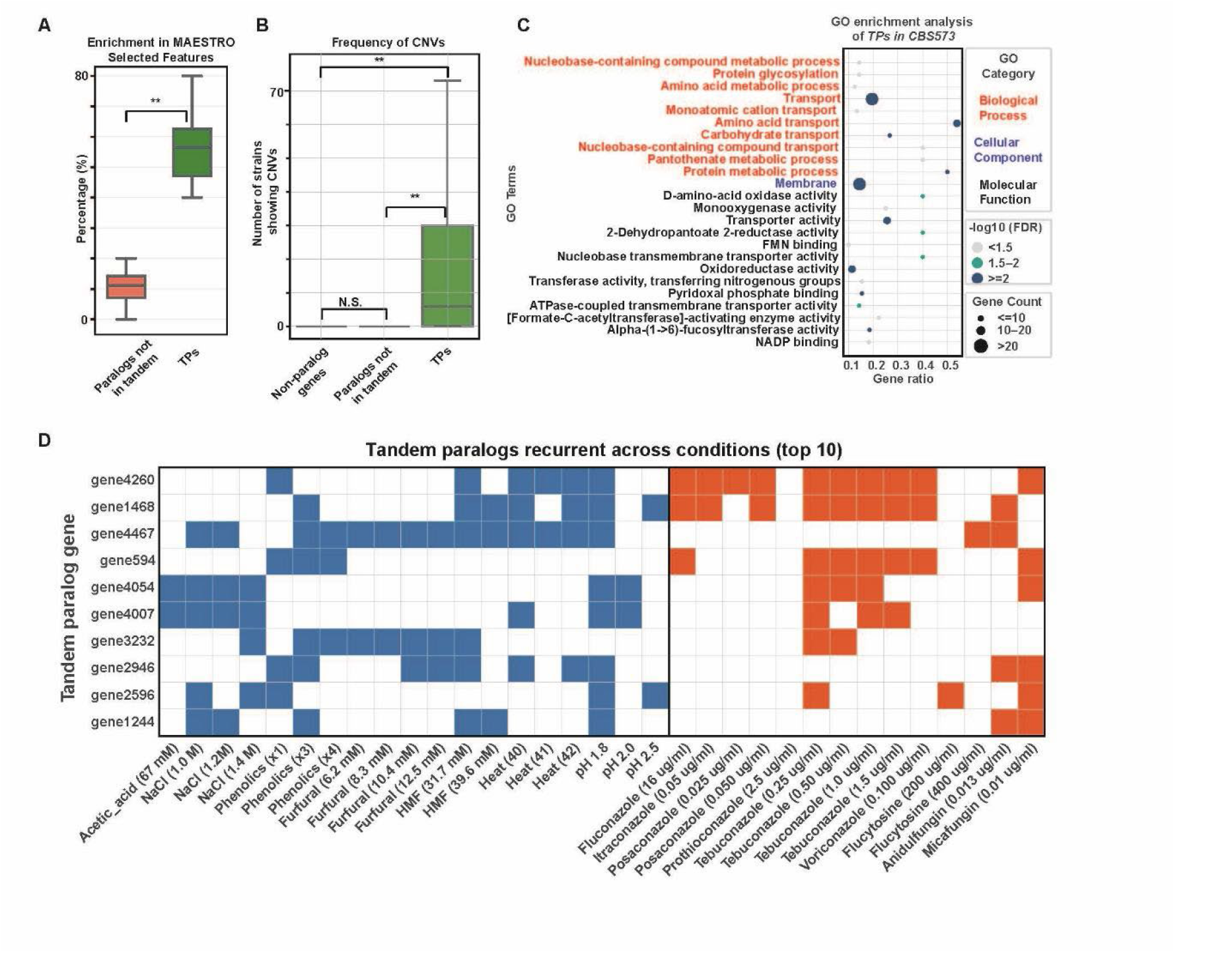
CNVs of TPs are preferentially selected by MAESTO pipeline and recurrently associated with multiple stress responses. (A) CNVs of TPs were significantly more enriched among MAESTRO-selected features compared with non-tandem paralogs (paired t-test, p < 0.01). (B) TPs exhibited significantly higher CNV frequencies across strains relative to non-paralog genes and non-tandem paralogs (Mann–Whitney U test, p < 0.01; N.S., not significant). (C) GO enrichment analysis of TPs in the CBS573 reference genome showed strong enrichment for transporters and membrane-associated functions. (D) Heatmap of the top 10 recurrently selected TPs across stress conditions, highlighting potentially pleiotropic candidates such as gene4260, which was consistently associated with multiple stressors.

### Pleiotropic roles of tandem paralogs gene4260/gene4261 in stress tolerance

To test whether gene4260 contributes to these traits, we set out to characterize loss-of-function mutants under the selected stress conditions. However, because tandem paralogs often display highly similar read-depth patterns (especially gene4260 and gene4261 share 99.04% sequence identity), their CNV features are highly collinear. In the MAESTRO pipeline, which uses LASSO regression as one of the feature selection methods, such collinearity can cause the model to arbitrarily retain one paralog while discarding the other. To address this, we generated a double mutant of the tandem paralogs gene4260/gene4261 rather than single mutants. This strategy also minimizes potential redundancy between the two genes and provides clearer phenotypic outcomes for functional analysis. For experimental validation, we selected IO021 as the benchmark strain due to its superior tolerance for lignocellulosic hydrolysates of energy crops (Fig. S24), a low-cost feedstock that supports the production of economically viable bioproducts in the bioeconomy.

As predicted, deletion of gene4260/gene4261 revealed pleiotropic effects across multiple stresses (Fig. 5A). In addition to its role in fluconazole responses, the double mutant gene4260Δ/gene4261Δ showed remarkably reduced growth under phenolics, HMF, and heat stress compared with the wild-type strain (WT). However, under low-pH conditions (pH 1.8), the mutant and WT strains grew similarly (Fig. 5B), contrary to expectations based on the association analysis. Interestingly, although biomass readouts from the BioLector microbioreactor for the gene4260Δ/gene4261Δ mutant appeared to be higher than WT during stationary phase, the final OD of the mutant was not higher than WT at the end point. The elevated biomass readings are likely due to altered cell properties in the mutant, such as increased aggregation or settling, rather than true increases in cell density. Although overall growth did not differ under this assay, it is possible that differences would become apparent under alternative measurement approaches that are less affected by aggregation.

**Figure 5.**
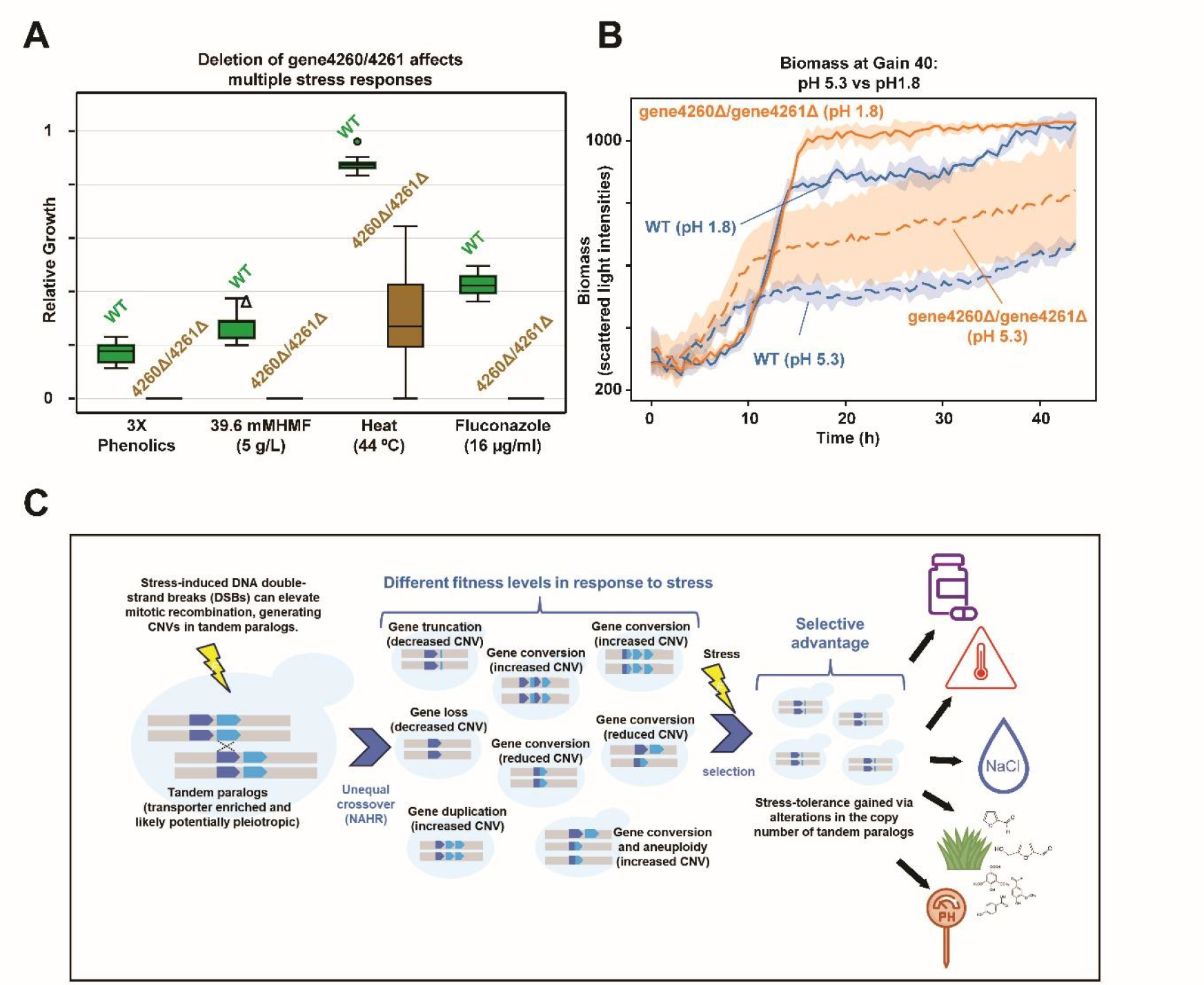
Pleiotropic roles of tandem paralogs gene4260/gene4261 in stress tolerance and evolutionary model of TP CNVs. (A) Growth phenotypes of WT and gene4260/gene4261Δ strains under selected stress conditions. The double mutant showed reduced tolerance to fluconazole, 3X phenolics (0.96 g/L vanillin, 0.66 g/L syringic acid, and 0.3 g/L 4-hydroxybenzoic acid), HMF, and heat compared with WT. (B) Biomass accumulation under low-pH conditions. WT and mutant strains grew similarly at pH 1.8, indicating that the predicted association of gene4260 CNVs with acid tolerance was not supported by knockout validation. (C) Model illustrating how CNVs and multifunctional tandem paralogs (TPs) dynamics contribute to adaptive plasticity. Based on their recurrent association with multiple stress responses, we hypothesize that NAHR-driven amplification or loss of TPs can shift dosage or generate novel functions through in-frame fusions. Such CNV dynamics could enable rapid adaptation to diverse stresses and promote the evolution of cross-resistance within the shared resistome network.

Collectively, these findings guided us to propose a TP CNV model that may represent a general strategy of fungal stress adaptation. In this model, TPs are recurrent CNV hotspots that undergo amplification, reduction, or recombination. Such CNV dynamics provide two adaptive advantages: (i) rapid, dosage-based tuning of stress-response genes, and (ii) opportunities for recombination between tandem copies that can generate chimeric alleles with novel functions. Together, these features make TPs highly flexible genetic modules that can drive multi-stress tolerance in *P. kudriavzevii* (Fig. 5C).

## DISCUSSION

While *S. cerevisiae* has long been a biotechnology workhorse, its sensitivity to industrial stress conditions increases production costs due to the need for detoxification and neutralization steps. In contrast, although non-model yeasts *P. kudriavzevii* can naturally tolerate multiple industrial-relevant stressors, they remain under-characterized, limiting their potential in industrial applications. At the same time, *P. kudriavzevii* has been known for its intrinsic resistance to fluconazole and was listed among the WHO’s fungal priority pathogens. Previous studies have shown that *P. kudriavzevii* can acquire drug resistance through overexpression of efflux pumps such as ABC1 and ABC2, and through mutations in the antifungal target gene ERG11. However, outside clinical settings where fluconazole is not present, it remains unclear what selection pressures drive environmental isolates to evolve drug resistance and give rise to resistant strains. The possibility that drug resistance can emerge outside clinical settings highlights a broader One Health concern, where environmental adaptation may inadvertently generate resistant strains. Understanding the genetic basis of stress tolerance in *P. kudriavzevii* is therefore critical not only for developing robust production strains but also for anticipating and mitigating biosafety risks.

In this study, we collected 170 isolates and developed an ML-assisted GWAS pipeline to dissect genotype–phenotype associations across both industrial and drug-related stressors. Because *P. kudriavzevii* is widespread across diverse habitats^25^ and frequently associated with plants outside of clinical settings^52^, environmental hotspots such as compost piles may serve as reservoirs where antifungal resistance emerges^4,53^. This raises a One Health concern that agricultural azole use could accelerate the emergence of azole-resistant *P. kudriavzevii* strains, as seen with drug-resistant *Aspergillus fumigatus*^*4*^. Indeed, we have found that the resistance to clinical fluconazole resistance is highly correlated with resistance to the agricultural triazole fungicide tebuconazole (Person correlation coefficient=0.71, p-value<0.01) among the strains. Moreover, fluconazole resistance is not significantly higher in clinical isolates than in environmental strains (Mann–Whitney test, p-value = 0.415), suggesting that environmental strains are already as drug-resistant as those from clinical settings. These patterns imply that agricultural fungicide use alone cannot account for all the observed resistance. Instead, our analyses reveal that many TPs are associated with both environmental stress tolerance and drug resistance (Fig. 4D). Such pleiotropic roles lead to an alternative evolutionary trajectory: resistance traits may have first evolved as adaptations to harsh environmental conditions and were later co-opted for antifungal resistance in clinical settings. For example, as seen with gene4260/gene4261, the same membrane transporters that confer fluconazole resistance also enhance tolerance to heat, HMF, and phenolics.

These observations are consistent with previous reports of cross-protection, where adaptation to one stressor unexpectedly enhances tolerance to others. A recent study showed that salt stress can increase high-temperature ethanol production, and one of the mechanisms involved up-regulation of alcohol dehydrogenase expression^54^. Supporting this result, we found that the alcohol dehydrogenase TP pair gene4467/gene4468 is associated with both NaCl and heat tolerance (Fig. 4D). In addition, adaptive laboratory evolution (ALE) of acetic acid-tolerant *P. kudriavzevii* strains was reported to exhibit enhanced multistress tolerance, including tolerance to heat, ethanol, osmotic stress, furfural, HMF, and vanillin^55^. Given that we identified the TP pair gene4260/gene4261 contributing to tolerance against heat, HMF, and vanillin (Fig. 5A), it is plausible that the acetic acid-tolerant ALE strains increased copy number of gene4260/gene4261. Other candidate TPs such as the aldo/keto reductase pair gene4054/gene4055 and the amino acid permease pair gene4006/gene4007 are associated with both acetic acid and NaCl tolerance (Fig. 4D), and thus represent strong targets for testing the genetic basis of cross-protection in ALE-derived strains. In this context, our TP CNV model suggests that ALE strategies intended to improve industrial traits may also incidentally expand drug resistance by co-opting multifunctional TP loci such as gene4260/gene4261.

Taken together, our findings extend current knowledge of gene CNVs and TPs in fungal adaptive evolution, highlighting CNV and TP dynamics as a general mechanism of fungal adaptive evolution (Fig. 5C). For instance, studies in Candida species have shown that CNVs contribute to antifungal resistance by amplifying drug target genes, often through tandem duplications^56–59^. Beyond the potential dosage changes via CNVs, homologous recombination within CNVs can generate in-frame fusions at TP loci, resulting in chimeric genes with altered functions. For example, expressing different chimeric ABC11-ABC1 TP alleles from *P. kudriavzevii* in *S. cerevisiae* leads to distinct antifungal sensitivities^60^. Similarly, in a population genomics study of *Candida parapsilosis*, the TP pair RTA3–RTA2 undergoes recurrent CNV and increased RTA3 copy number strongly correlates with resistance to miltefosine^18^. Although most of *C. parapsilosis* strains in the study were human-associated, RTA3 amplifications were also found in environmental isolates and appear to arise independently of drug exposure^61^. This supports our model in which TP CNVs enable adaptation to environmental stressors that can incidentally produce antifungal resistance.

More broadly, recent studies in *Candida albicans* have shown that long repeat sequences serve as hotspots for CNV, LOH, and chromosomal inversions^62^, and TPs in *P. kudriavzevii* can be viewed as localized repeat sequences that act as hotspots for CNV and recombination. Moreover, expandable and reversible CNV amplifications have been shown to drive rapid antifungal adaptation in *C. albicans*^*22*^, supporting our hypothesis that TP CNVs may act as flexible, repeat-like elements that can be expandable and reversible. These findings reinforce our conclusion that the TP CNV model uncovered in *P. kudriavzevii* reflects a broader evolutionary strategy for rapid fungal adaptation (Fig. 5C).

In studies of uncommon and rare diseases, sample sizes are inherently limited, which is similar to our situation in studying non-model yeasts. While expanding the cohort size would naturally improve predictive accuracy, we also recognize that not every study can achieve the scale needed for classical GWAS. Our results show that even with a limited dataset of 170 strains, MAESTRO can consistently identify features that are biologically meaningful. These findings reveal tandem paralogs and CNV dynamics as key mechanisms of fungal stress adaptation, while also demonstrating MAESTRO as a practical framework for extracting interpretable associations when large-scale datasets are not feasible. Beyond non-model yeasts, MAESTRO may be broadly useful in other contexts where sample size is constrained, such as non-model plant and microbial populations, ALE lines, conservation genetics of endangered species, and small clinical cohorts.

## ACKNOWLEDGEMENTS

This work was supported by the US Department of Energy Center for Advanced Bioenergy and Bioproducts Innovation (DOE, Office of Science contract DE-SC0018420 and DE-AC02-05CH1123) and Biosystems Design program (DOE, Office of Science contract DE-SC0018260 and DE-AC02-05CH1123). The work (proposal: 10.46936/10.25585/60001019) conducted by the US Department of Energy Joint Genome Institute (https://ror.org/04xm1d337), a DOE Office of Science User Facility, is supported by the Office of Science of the US Department of Energy operated under Contract No. DE-AC02-05CH11231. Any opinions, findings, conclusions, or recommendations expressed in this publication are those of the author(s) and do not necessarily reflect the views of DOE. Y.-P.L. is supported by the Taiwan NSTC Young Scholar Fellowship Einstein Program (112-2636-E-002-005). Research in the Hittinger Lab is supported by the US National Science Foundation under Grant No. DEB-2110403, the USDA National Institute of Food and Agriculture (Hatch Projects 1020204 and 7005101), in part by the DOE Great Lakes Bioenergy Research Center (DOE BER Office of Science DE– SC0018409, and an H. I. Romnes Faculty Fellowship, supported by the Office of the Vice Chancellor for Research and Graduate Education with funding from the Wisconsin Alumni Research Foundation. We thank Anita Wahler for professional editing of this manuscript. We are grateful that the US Department of Agriculture, Agricultural Research Service Culture Collection (Northern Regional Research Laboratory [NRRL]) Database provided us with nearly 60 strains free of charge.

## AUTHOR CONTRIBUTIONS

Y.Y., P.-H.H., and Y.S. conceived of the research project. P.-H.H. and Z.Z. were responsible for preparing the 162 samples for genomic sequencing. Another eight strains for genome sequencing were prepared by D.A.O. and C.T.H. Genomic analysis was conducted by P.-H.H. Phenolic analysis was conducted by Y.S.. A.S.S. performed the maximum-likelihood phylogenetic tree and GO analyses. J.K. carried out the fermentation experiments for the benchmark strain screening, and B.D. provided the switchgrass hydrolysates. S.H. performed the draft genome analysis of the new benchmark strain. P.-H.H and S.D. performed the linear-mixed-model GWAS analysis. Genome engineering and verification experiments were performed by P.-H.H., Z.-Y.W., Y.S., and Z.F. MAESTRO was conceived of and implemented by P.-H.H., C.-I.Y., and Y.-P.L. P.-H.H., Y.S., and Y.Y. drafted the manuscript and generated figures and tables. All authors participated in reviewing, editing, and approving the final version.

